# Relative quantification of *Porphyromonas gingivalis*, *Treponema denticola*, *Tannerella forsythia* and *Aggregatibacter actinomycetemcomitans* high-risk bacterial species in Romanian patients evaluated for periodontal disease

**DOI:** 10.1101/184770

**Authors:** Cosac Ion Constantin, Ionica Consuel, Ratiu Attila Cristian, Savu Lorand

## Abstract

Three bacterial species pertaining to the red complex (*Porphyromonas gingivalis, Treponema denticola,* and *Tannerella forsythia*) and *Aggregatibacter actinomycetemcomitans* were investigated in relation to the incidence and severity of periodontal disease. A total of 259 patients were included in this study, 179 being diagnosed with periodontal disease. The gingival crevicular fluid samples were obtained from periodontal pockets and the presence and levels of target bacteria were assessed following DNA extraction and real-time quantitative PCR. Our results account for significant positive associations between the number of bacterial species from the red complex coexisting within a patient and several clinical signs (gingival bleeding, inflammation and bone deterioration). A similar positive association was found between bacterial load of the red complex species and the clinical Case Type diagnostic of the periodontal disease, as well as the probing depth with the most evident results for *T. denticola.* In conclusion, our study, a first for the Romanian population, confirms previous results found elsewhere and finds a possible regional pathogenic specificity for *T. denticola* as a major factor for periodontitis severity.

## Introduction

Periodontal disease is a multifactorial affliction that is becoming increasingly common in middle-aged people, mainly due to the lack of attention to distinctive clinical signs (Chen *et al,* 2005). Genetic predisposition, lifestyle (smoking, stress, and oral hygiene), as well as socio-economic status, precede and modulate the development of the bacterial biofilm present in the gingival sulcus, which represents the main cause of gingivitis and periodontal disease (Mariotti and Arthur, 2015; Chen *et al,* 2005).

Periodontal disease is commonly recognized as the association between gingival inflammation and the pathological detachment of the connective tissue (Armitage, 1995). Gingival inflammation is mainly caused by infections with specific groups of oral bacteria (Armitage, 2003). The treatment of periodontal disease consists in both surgical (gum surgery and bone grafts) and non-surgical approaches (removal of the microbial plaque and administration of antibiotics). Identification of pathogenic species of bacteria is essential for the antibiotic treatment (Loloya-Rodriguez *et al.,* 2014). In addition, quantifying the bacterial load can help monitoring the progression of the treatment.

Several methodologies were used to investigate the periodontal pathogenic bacterial load for therapy purposes, such as targeting 16S RNA genes (Armitage, 2003). According to the previous clinical assessments, periodontal pathogenic bacteria are usually grouped into three main classes (or complexes): red, orange and green. The Gram-negative anaerobic bacteria in the red complex (*P. gingivalis, T. denticola, T. forsythia*), representing the most aggressive class, along with *A. actinomycetemcomitans* from the green complex, are believed to be major etiologic agents for determining the severity of periodontal disease (Decat *et al.,* 2012; The Research, Science and Therapy Committee of the American Academy of Periodontology, 1999; Tettamanti *et al.,* 2017). Bacteria complexes, other than the red one, are also important but have a lesser impact on the severity of the disease. It appears that orange and green complexes are associated with long term gingivitis, which precedes the actual periodontal disease, while the red complex are the late colonizers of the plaque biofilm (Teles *et al.,* 2013). *A. actinomycetemcomitans* has been found to be one of the first and most important etiological agents that invades the subgingival plaque and set up the environment for other bacterial species (Mariotti and Arthur, 2015).

Several studies regarding the prevalence and bacterial load of various species of pathogenic bacteria found differences among the focus populations (Tettamanti *et al.,* 2017). These results are relevant in order to successfully elaborate strategies using specific antibiotics for targeting the most frequent/aggressive bacterial strains in a variety of populations.

Our study represents the first assessment of the pathogenic bacteria responsible for periodontal diseases in Romanian patients, its main objective being to assess the correspondence between the presence and load of the red complex bacteria and respectively *A. actinomycetemcomitans* with various levels of periodontal disease.

## Materials and methods

### Subject population

This study included 259 periodontally untreated patients with an average age of 44 years (sd ± 11), 51% of them being women. Additional information on evaluated patients included self-recorded categories of age, sex, and smoker status, as well as rarely stated pregnancy and relevant medical conditions such as diabetes, immunodepression, heart diseases or hereditary periodontal disease. Supplementary data provided by the dental clinic included description of symptoms (gingival bleeding, redness, inflammation, bone deterioration, and halitosis), clinical degree of periodontal disease (Case Type I - gingivitis, Case Type II - mild periodontitis, Case Type III - moderate periodontitis, Case Type IV - advanced periodontitis, and Case Type V - refractory periodontitis) (Armitage, 2003), and depth of pocket probing (range of 1 mm to 15 mm).

### Sample collection

Samples of gingival crevicular fluid were collected at the dental clinic using five sterile paper tips, each inserted into a different periodontal pocket and held there for approximately 10 seconds. Afterwards, the paper tips were transferred to a single sterile tube and sent to our laboratory for DNA extraction and bacterial quantification. Therefore, total bacterial load represented a pool of different species present in the tested periodontal pockets.

### Bacterial DNA extraction

Paper tips were resuspended overnight in 1 mL of 1.5X PBS solution. Bacterial pellets were obtained after 10 minutes of centrifugation at 16,000 g and, subsequently, 200 μL of PBS containing the resuspended bacteria were further processed using the Viral Kit I (Favorgene) according to manufacturer’s instructions. Briefly, samples were supplemented with 200 μL of PBS and 570 μL of VNE buffer (enriched with DNA carrier) and, after 10 minutes of incubation at room temperature, 570 μL absolute ethanol were added. Subsequently, the resulting solutions were loaded on individual DNA separation columns that were washed one time with 500 μL wash buffer I and two times with 750 μL wash buffer II. Finally, DNA was eluted in 80 μL pre-heated nuclease-free water.

### Real-time quantitative PCR (qRT-PCR)

Bacterial quantification was performed on a LightCycler 2.0 machine (Roche). Single PCRs targeting individual bacterial species were performed in a total volume of 20 μL containing 1.25X colourless buffer (Promega), 0.25 mM dNTPs, 4 mM MgCl2, 1.5 units of PQ Taq (Promega), 1 μL of 1X SYBR Green I, 0.5 mM of each primer (primer sequences are listed in table 1) and 5 μL of DNA (Kirakodu *et al.,* 2008; Masunaga *et al.,* 2010; Bastos *et* al.,2011).

**Table 1.**
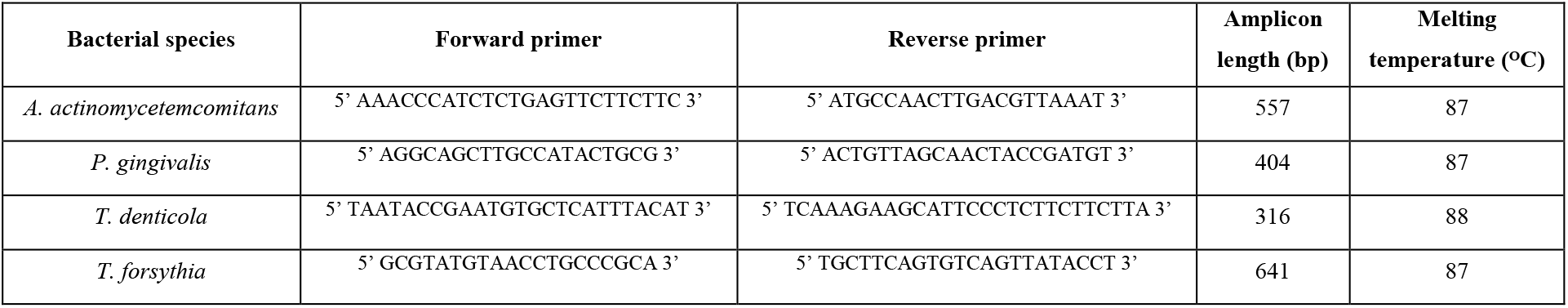
Primer sequences used for PCR amplifications.

The amplification comprised of an initial 5 minutes denaturation at 95°C, 40 cycles consisting in 10 seconds at 95°C, 5 seconds at 55°C, and 25 seconds at 72°C, followed by High Resolution Melt (HRM) analysis in order to monitor the species-specific melting point.

The relative quantitation of bacterial load was achieved by comparison with a standard amplification curve obtained starting from genomic DNA corresponding to 6.0E6 colony founding units (CFUs) of *P. gingivalis* isolated from pure cultures.

The LightCycler 1.5 software was used to infer the cycle threshold (Ct) values and to perform the HRM analysis.

### Statistical analysis

We applied chi-square analysis to test for association between the number of red-complex species present per patient and presence of each symptom. To assess each species’ load as a function of probing depth we employed Pearson correlation. The same test was used to evaluate the association of pairs of species. Finally, a Kruskal-Wallis non-parametric ANOVA was applied separately to each species’ bacterial load to test for the association with the severity of periodontal disease as clinically diagnosed by dentists (i.e. Case Type). All tests were performed using the Minitab v.16 software or R software environment using a 5% level of significance, two tailed.

## Results and Discussions

In our approach, each bacterial species was targeted by individual qRT-PCR reactions containing specific primer pairs. The LightCycler 1.5 software computed the Ct values that were compared with the Ct inferred from the amplification curve of a standard sample containing genomic DNA equivalent to 6E6 CFUs of *P. gingivalis.* The target amplicons corresponding to specific bacterial species have different lengths and this feature could impact the amplification efficiency, due to their gincreased size that enhances the assay sensitivity (Peirson *et al.,* 2003). However, the fact that Ct values are estimated in the early cycles of exponential amplification, and considering that errors of up to few hundred percent may be tolerated when trends or relative big changes in amounts are measured (Bar *et al.,* 2012), PCR efficiencies are adequately comparable for the study’s purpose.

Bacterial load of all species ranged between 1.0E6 and 1.0E14 CFUs (table 2). Samples were positive for at least one (N = 10, 3.86% of cases), two (N = 42, 16.22% of cases), three (N = 151, 58.30% of cases) and all four (N = 49, 18.92% of cases) investigated pathogenic bacteria, with varying degrees of abundance across species. Several samples (N = 155, 59.8% of cases) were also positive for additional bacteria pertaining to orange and green complexes (data not shown).

**Table 2.** Primer sequences used for PCR amplifications.

| Bacterial species | Mean CFUs | Standard deviation | Median CFUs |
| --- | --- | --- | --- |
| *A. actinomycetemcomitans* | E9.87 | E1.98 | E10 |
| *P. gingivalis* | E12.9 | E1.83 | E14 |
| *T. denticola* | E12 | E1.96 | E12 |
| *T. forsythia* | E12.6 | E1.71 | E13 |

*A. actinomycetemcomitans* was by far the least present species (N = 61, 24% of cases), while the other bacterial species were detected more frequently: *T. denticola* (N = 219, 85% of cases), *T. forsythia* (N = 227, 88% of cases), and *P. gingivalis* (N = 236, 91% of cases).

Student’s t-test and regression analysis showed that there are no associations between bacterial loads and patients’ gender or age, results consistent with those provided by other studies (Farias *et al.,* 2012).

Evaluation of the bacterial community comprising the analysed species indicated significant positive association between *T. denticola* and *T. forsythia* (r = 0.589, p < 0.001), *P. gingivalis* and *T. denticola* (r = 0.437, p < 0.001), *P. gingivalis* and *T. forsythia* (r = 0.470, p < 0.001), and *A. actinomycetemcomitans* and *T. forsythia* (r = 0.354, p = 0.006), when tested as bacterial load using Pearson correlation. In contrast, we did not find significant associations between *T. denticola* and *A. actinomycetemcomitans* (r = -0.058, p = 0.674), and respectively *P. gingivalis* and *A. actinomycetemcomitans* (r = 0.010, p = 0.945). The latter situation, illustrating a seeming conflict between the involved bacterial strains, could be explained by the fact that the major virulence factors of *P. gingivalis* consisting in the extracellular proteinases called Lys-gingipains (Gorman *et al.,* 2015) are involved in the detachment and decrease of *A. actinomycetemcomitans* biofilms (Haraguchi *et al.,* 2014). The *P. gingivalis* gingipains seem to also have positive effects on *T. forsythia* growth and are crucial for its co-adhesion with *T. denticola* (Bao *et al.,* 2014) thus explaining their strong association at the same lesion. A previous study investigating the grouping of bacterial species existing in subgingival plaque found that the most tightly related group, forming a major complex related to worsened pocket depth and gingival bleeding, was composed of *P. gingivalis, T. denticola* and *T. forsythia* (Socransky *et al.,* 1998). Interestingly, the surprising significant association between *A. actinomycetemcomitans* and *T. forsythia* could be in fact the consequence of the fact that the latter uses other virulence mechanisms instead of Lys-gingipains (Friedrich *et al.,* 2015), thus probably not hindering the development of *A. actinomycetemcomitans*.

We tested for associations between the presence or absence of various clinical symptoms scored for patients and the number of bacterial species pertaining to the red complex. Some of the cases exhibited more than one symptom, thus they were counted each time an individual symptom was considered. Chi-square analysis of data representing the number of cases without or with symptoms when none, one, two, or all three red complex species were present, did not yield significant trends when redness or halitosis were analysed, but showed strong statistical significance (p < 10^-3^) for gingival bleeding, inflammation and bone deterioration. These symptoms were exceptionally common in cases displaying three bacterial species, more precisely, 167, 124 and 147 patients presenting gingival bleeding, inflammation or bone deterioration, respectively (figure 1). Similar results were previously reported when coexistence of all three species from the red complex determined increases in maxillary and mandibular alveolar bone resorption (Suzuki *et al.,* 2013) or periodontal disease severity (Lanza *et al.,* 2016).

**Figure 1.**
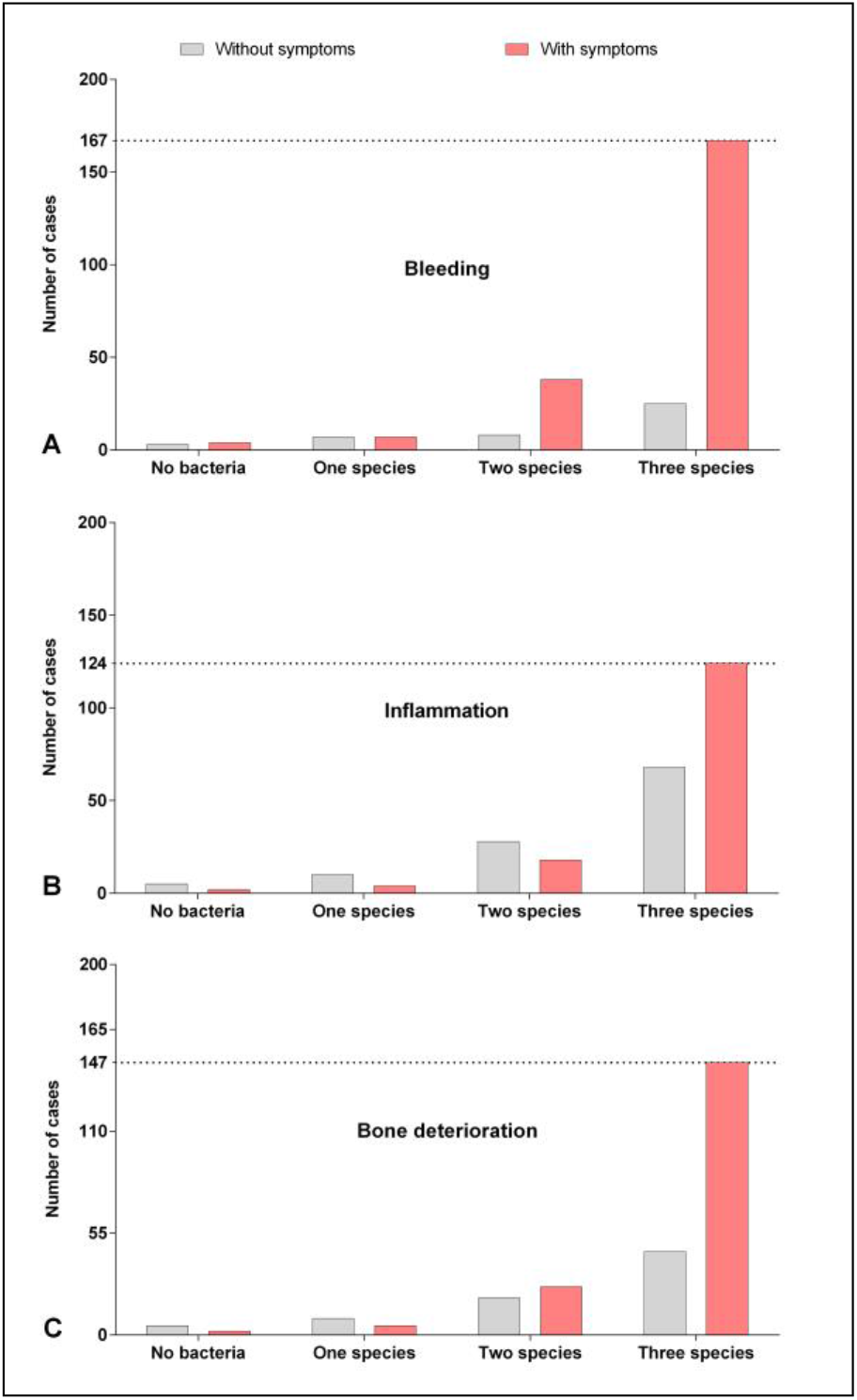
Associations between the number of bacterial species identified in a given patient and the presence of gingival bleeding (Figure 1A), inflammation (Figure 1B) and bone deterioration (Figure 1C) clinical symptoms. For each symptom, we emphasized the maximum number of cases presenting the symptoms, which is strongly correlated with the influences exerted concomitantly by the various bacterial species from the red complex.

A stepwise regression analysis on cases with all red complex bacteria present (N = 130) retained only the load of *T. denticola* as significant for the maximum probing depth (t = 4.47, p < 10^-3^, R-sq adj. = 12.84%, Mallows Cp = 2.2). When each bacterial species was individually tested (Pearson correlation), we were able to show a decrease of *A. actinomycetemcomitans* bacterial load with the maximum depth of pocket probing, although the result was not statistical significant (r = -0.241, p = 0.133). This result requires further investigation, given the facultative anaerobic nature of this species (Tanner, 2015), as well as the competitive relationship between *P. gingivalis* and *A. actinomycetemcomitans.* (Haraguchi *et al.,* 2014). In contrast, red complex bacterial load appeared to be positively correlated with maximum probing depth, significant associations being calculated for *P. gingivalis* (r = 0.238, p = 0.003) and *T. denticola* (r = 0.361, p < 0.001), but not for *T. forsythia* (r = 0.111, p = 0.161). A study focused on adult Brazilians demonstrated the significant prevalence of *P. gingivalis* in deep pockets, but also of *T. forsythia* (Farias *et al.,* 2012). The fact that we identified *T. denticola* as bearing the strongest correlation with maximum probing depth could be representing a local trait, as several studies indicated that periodontal pathogens’ distribution and prevalence vary with geographic location (Tettamanti *et al.,* 2017) and ethnicity (Gatto *et al.,* 2014). Furthermore, the latter team found that in Italian population the strongest association with pocket depth is that of *T. forsythia,* which supports local bacterial specificity hypothesis.

A Kruskal-Wallis non-parametric ANOVA was applied to the bacterial load, separately for each red complex species, as a function of Case Type clinical diagnosis. Only 177 subjects were classified according to this criterion. An increase was detected for all three species from Case Type 1 to Case Type 5 (figure 2), which proved significant for *T. denticola* (H = 17.89, p = 0.001) and *P. gingivalis* (H = 13.87, p = 0.01), but not for *T. forsythia* (H = 8.14, p = 0.08). The Kruskal-Wallis test was chosen for its ability to control for non-normal data by analysing the ranks instead of actual bacterial load values, an aspect important for our data, which clustered in the upper part of the scale and was under-represented in the extreme Case Types 1 and 5. The results indicate an increase of the bacterial load along with the severity of diagnosis, although significant differences between levels were not always detected in a post-hoc evaluation. However, rather than strictly evaluating differences between each Case Type class, the purpose of the test was to assess the pattern of the relationship, if any. Such a tendency was also demonstrated in a Thai population (Wara-aswapati *et al.,* 2009) for the red complex bacteria. Since our analysis included patients with no bacterial load, we repeated the test without these null figures. The results remained significant only for *T. denticola* (H = 14.68, p = 0.005, N=156), but not for *P. gingivalis* (H = 5.60, p = 0.23, N=155) or *T. forsythia* (H = 5.87, p = 0.20, N=166). Compared to the red complex species, *A. actinomycetemcomitans* bacterial load showed no trend in relationship with the dental pathology diagnosis (H = 3.40, p = 0.33, N=43).

**Fig. 2.**
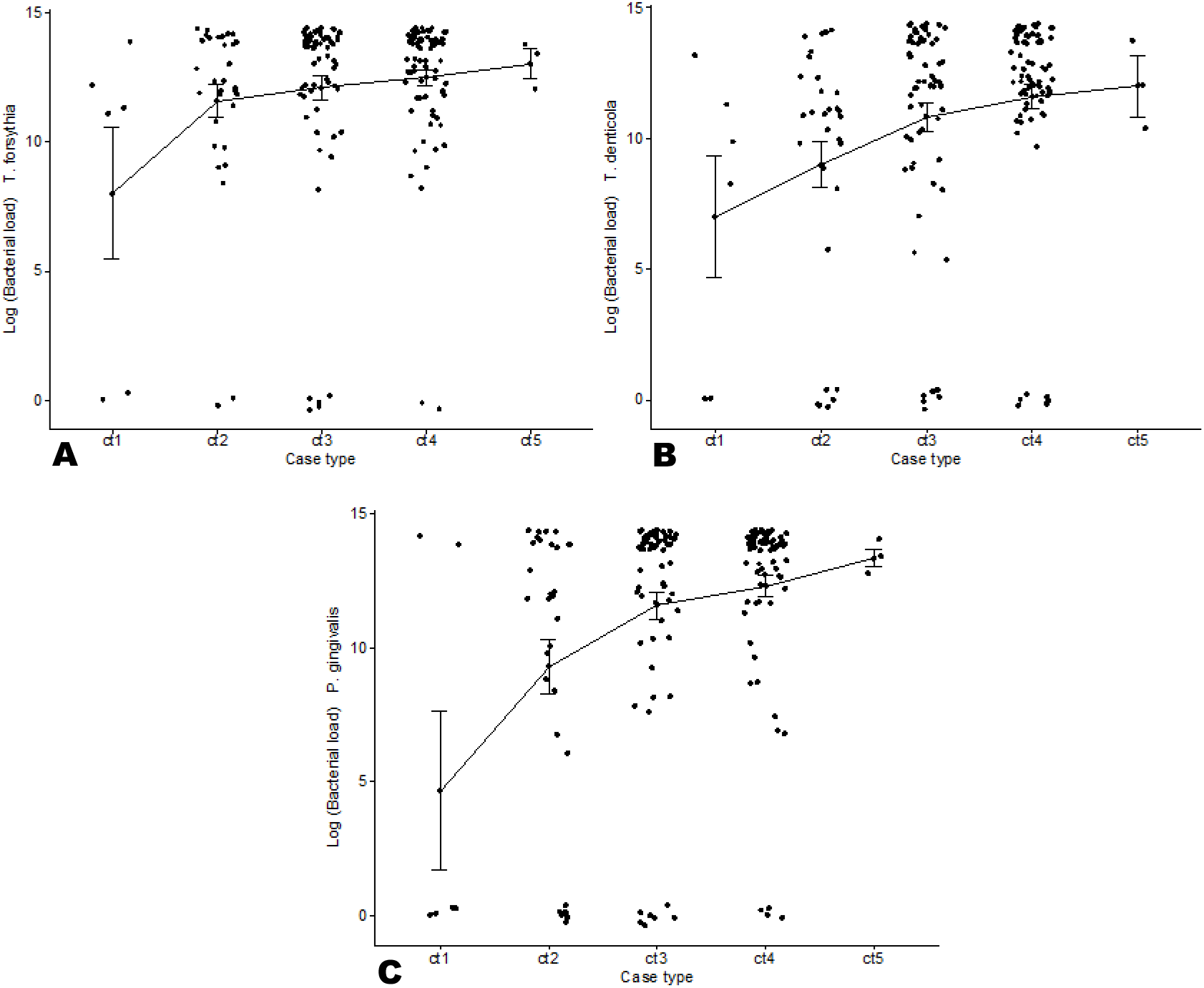
Graphical representation of logarithmic distribution of bacterial load (CFUs expressed as logarithmated values) as a function of Case Type diagnosis (ct1 to ct5) for *T. forsythia* (A), *T. denticola* (B) and *P. gingivalis* (C).

## Conclusion

In our study population, coexistence of red complex bacteria is positively associated with several dental pathologies, i.e. inflammation, gingival bleeding and bone deterioration, but not with halitosis or redness. No similar association was found for *A. actinomycetemcomitans*. Bacterial load did not associate as strongly with pathology, except for *P. gingivalis* and *T. denticola*, with regard to the depth of probing and the severity of periodontal disease as clinically diagnosed (Case Type 1 to 5). *P. gingivalis* and *T. denticola*, *P. gingivalis* and *T. forsythia*, *T. denticola* and *T. forsythia*, as well as *A. actinomycetemcomitans* and *T. forsythia* appear to associate with each other. In contrast to red complex bacteria, *A. actinomycetemcomitans* was identified in relatively few cases and apparently its bacterial load decreases with the depth of dental pockets, a result that needs to be eventually validated on a larger population sample.

Both *P. gingivalis* and *T. denticola,* but especially the latter, characterize dental damage in Romanian patients. The highest prevalence of *T. denticola* in severe cases of periodontal disease as well as its strongest positive correlation with maximum pocket depth could be representative for the Romanian population, a hypothesis that needs to be rigorously tested.

